# Clustering of colon, lung, and other cancer susceptibility genes with protein tyrosine phosphatases and protein kinases in multiple short genomic regions

**DOI:** 10.1101/2023.11.07.566108

**Authors:** Lei Quan, Peter Demant

## Abstract

Interactions of large gene families are poorly understood. We found that human, mouse, and rat colon and lung cancer susceptibility genes, presently considered as separate gene families, were frequently pairwise linked. The orthologous mouse map positions of 142 of 159 early discovered colon and lung cancer susceptibility genes formed 41 genomic clusters conserved >70 million years. These linked gene pairs concordantly affected both tumors and their majority was linked with two other gene families - protein tyrosine phosphatases and cancer driver protein kinases. 25% of both protein tyrosine phosphatases and protein kinases mapped <1 cM from a colon or lung cancer susceptibility gene, and 50% in <3 cM. Similar linkage was detected with most other human susceptibility genes that controlled 29 different cancer types. This concentration of tumor susceptibility genes with protein tyrosine phosphatases and driver protein kinases in multiple relatively short genomic regions suggests their possible functional diversity.

## Introduction

Common non-familial cancers were found in experimental animals to be controlled by multiple susceptibility genes with limited effect^1–3^ and many such loci were subsequently mapped in human genome wide association studies (GWAS)^4^. Until now, however, a systematic analysis of common features of genetics of colon and lung cancer susceptibility loci has not been reported and their genomic distribution was generally considered to be random. We present here comprehensive data demonstrating their pair-wise linkage in the genome, conserved between humans and mice for > 70 million years. In addition, these cancer susceptibility genes are linked to protein tyrosine phosphatases and driver protein kinases. These two types of enzymes were linked also to 29 other types of human cancer susceptibility loci.

## Results

### The common characteristics of colon and lung cancer susceptibility genes

#### a. Co-mapping of colon and lung cancer susceptibility loci

We have mapped in mice numerous susceptibility genes for colon and lung cancer using recombinant congenic (RC) strains^3^. 12 of the 15 mouse colon cancer susceptibility loci (Scc - Susceptibility to colon cancer) and 22 lung cancer susceptibility loci (mostly used symbols Sluc - Susceptibility to lung cancer or Pas - Pulmonary adenoma susceptibility) were frequently linked pair-wise together^5^, forming 19 clusters (14 of them shorter than 2.5 cM) that included also 10 pre-GWAS human and 3 rat^6^ colon cancer susceptibility loci.

This clustering was obvious in spite of different species, different tumor induction protocols, different strains^3.5,7^, and use of different carcinogens: 1,2 dimethylhydrazine (DMH), or azoxymethane (AOM)^5,7,8^ for colon cancers and N-ethyl-N-nitroso-urea (ENU), or ethyl-carbamate (urethane)^5,9^ for lung cancers.

#### b. GWAS data confirmed the intra- and inter-species co-localization of colon and lung cancer susceptibility loci

The first comprehensive GWAS data^10^ using the standard p-value threshold of 5 x 10-^8^ revealed 31 novel colon cancer susceptibility loci in humans^4^. Orthologous positions of 23 of these loci on the mouse chromosome map were in vicinity of mouse Sluc loci and confirmed our previous report of conserved co-localization of colon and lung cancer susceptibility^5^. With the previous data this revealed 27 clusters of paired colon-lung cancer susceptibility loci from human, mouse and rat ^11^, nine of which were < 2.0 cM long.

Additional GWAS^12,13^ detected 33 new human colon cancer susceptibility loci. Combining GWAS SNPs and Oncotarget markers detected 18 new human lung cancer susceptibility loci in populations of European origin^14^, and 5 lung cancer susceptibility loci in populations of Asian descent^15^, all at p <5 x 10^-8^. All these 23 new human lung cancer susceptibility loci were clustered with colon cancer susceptibility loci. Moreover, the lung cancer susceptibility gene VTlI1A^15^ was independently identified as a colon cancer susceptibility gene^16^. These data are summarized in Table 1.

**Table 1.**
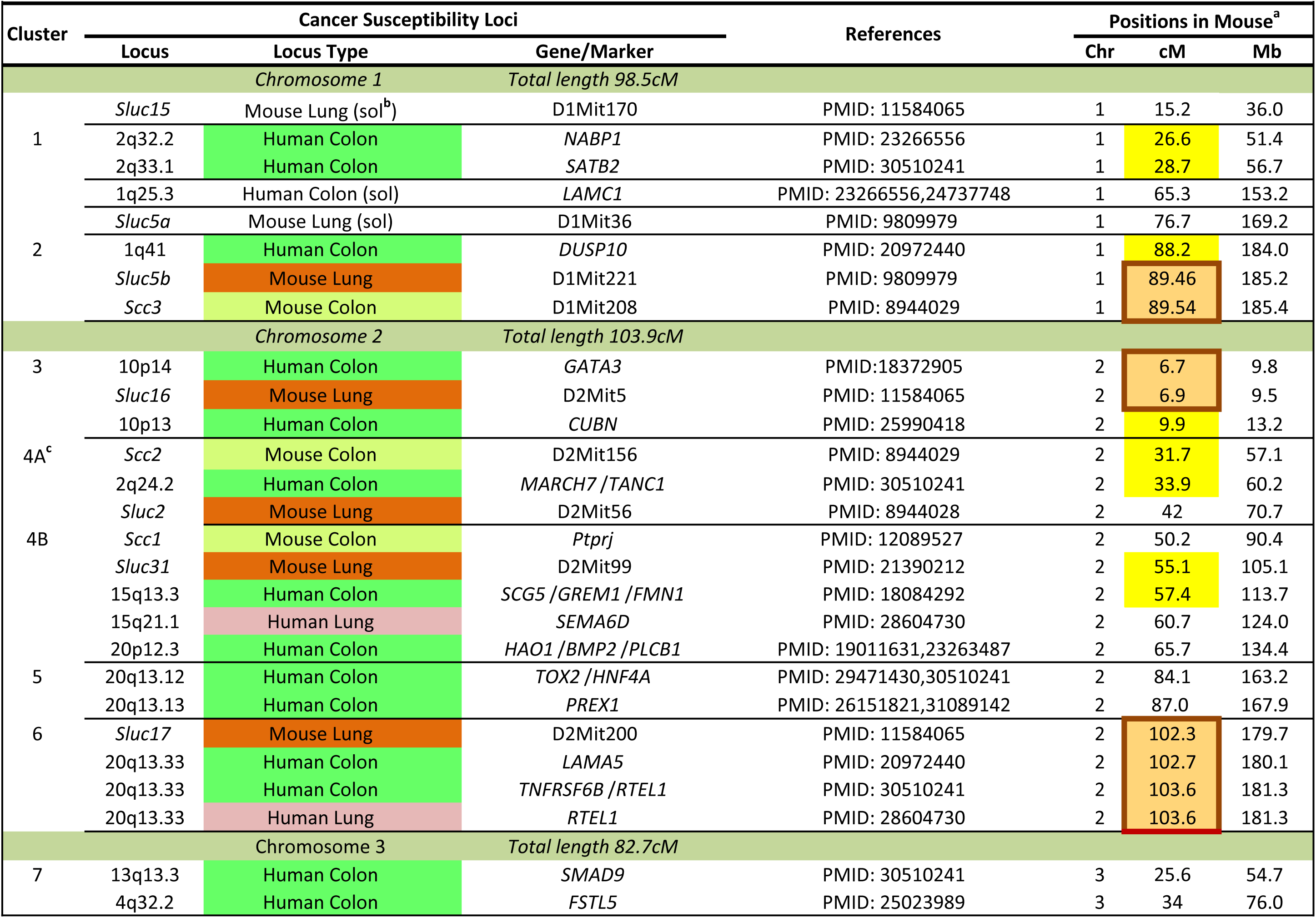
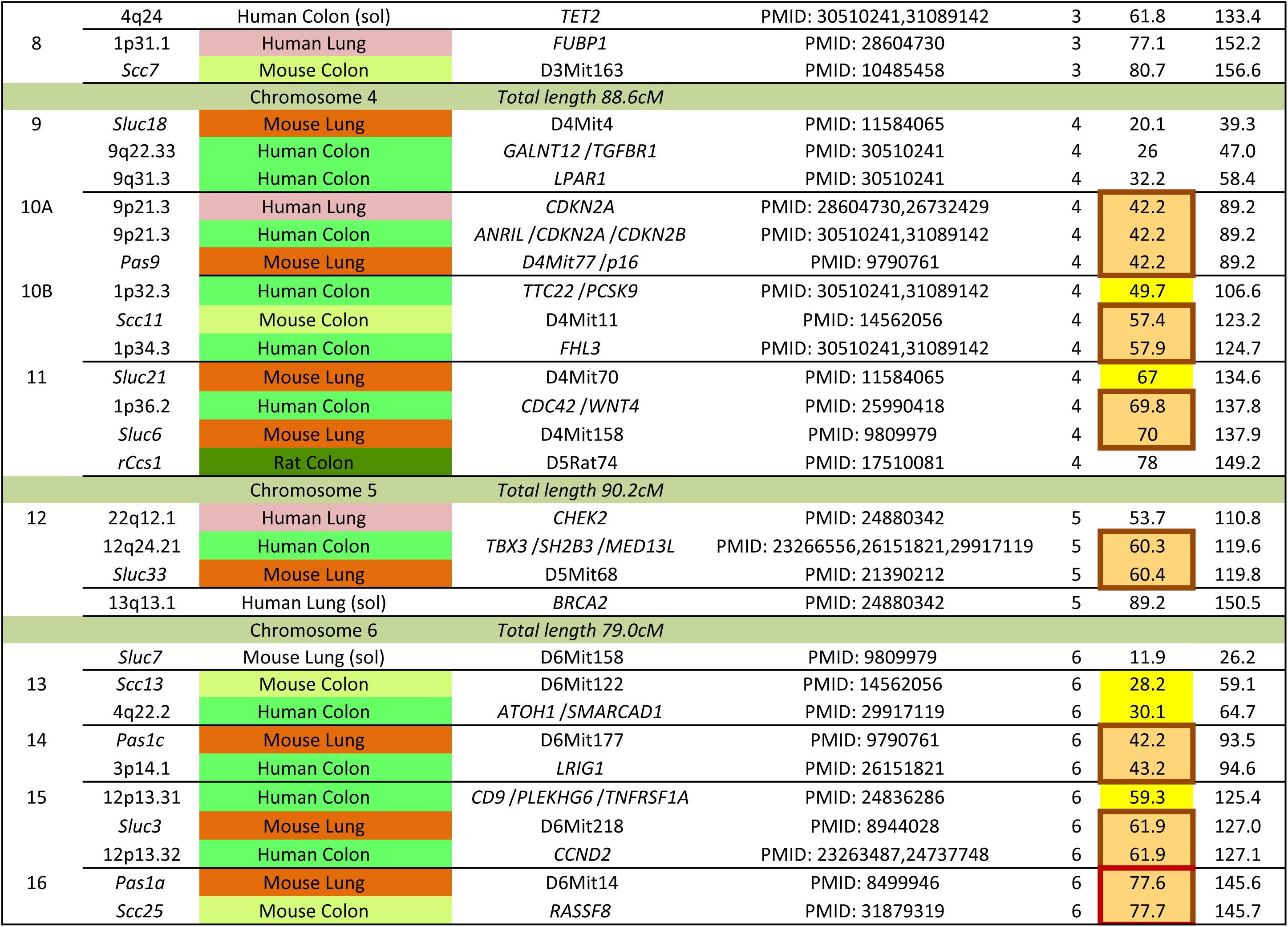
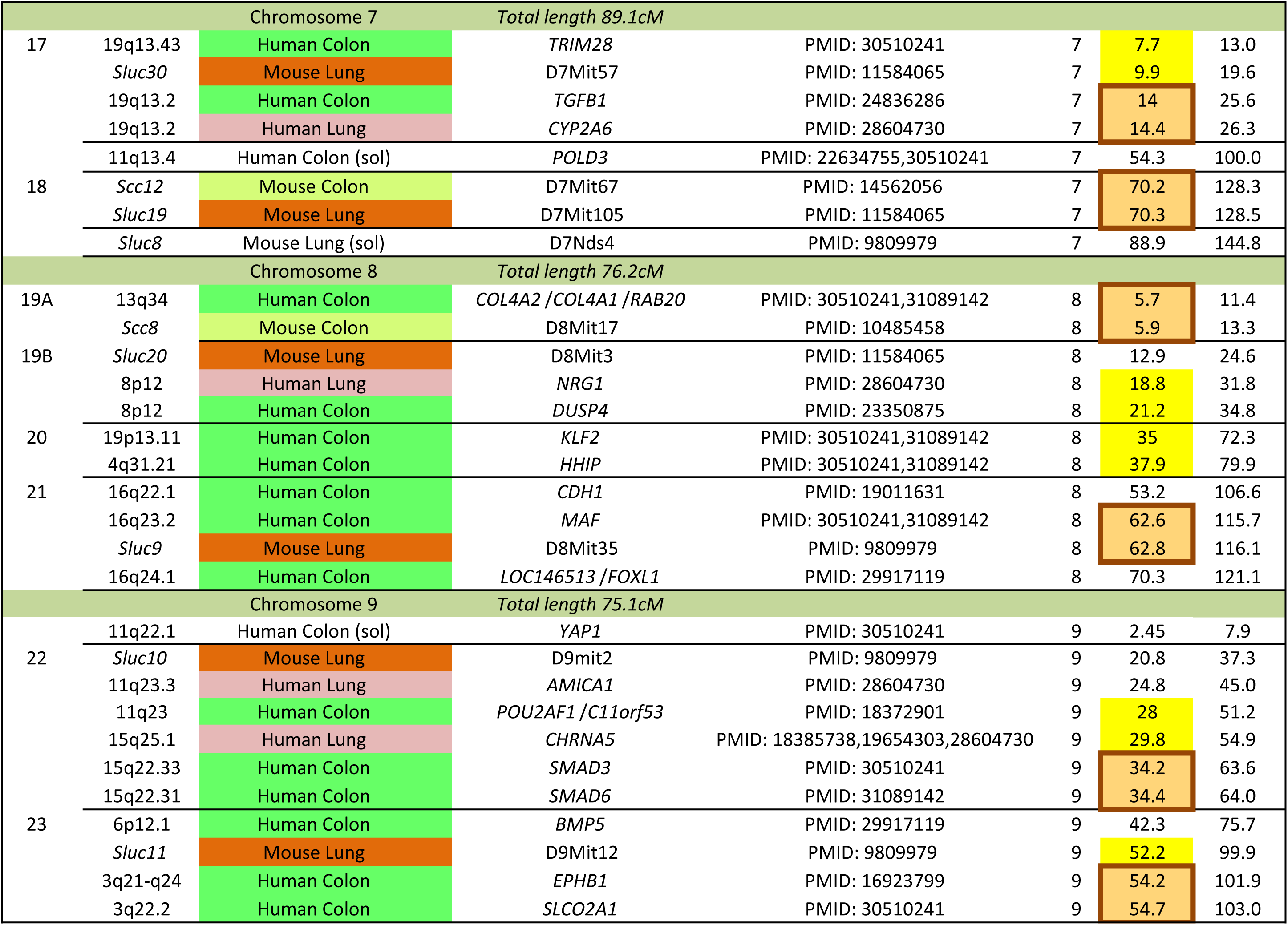
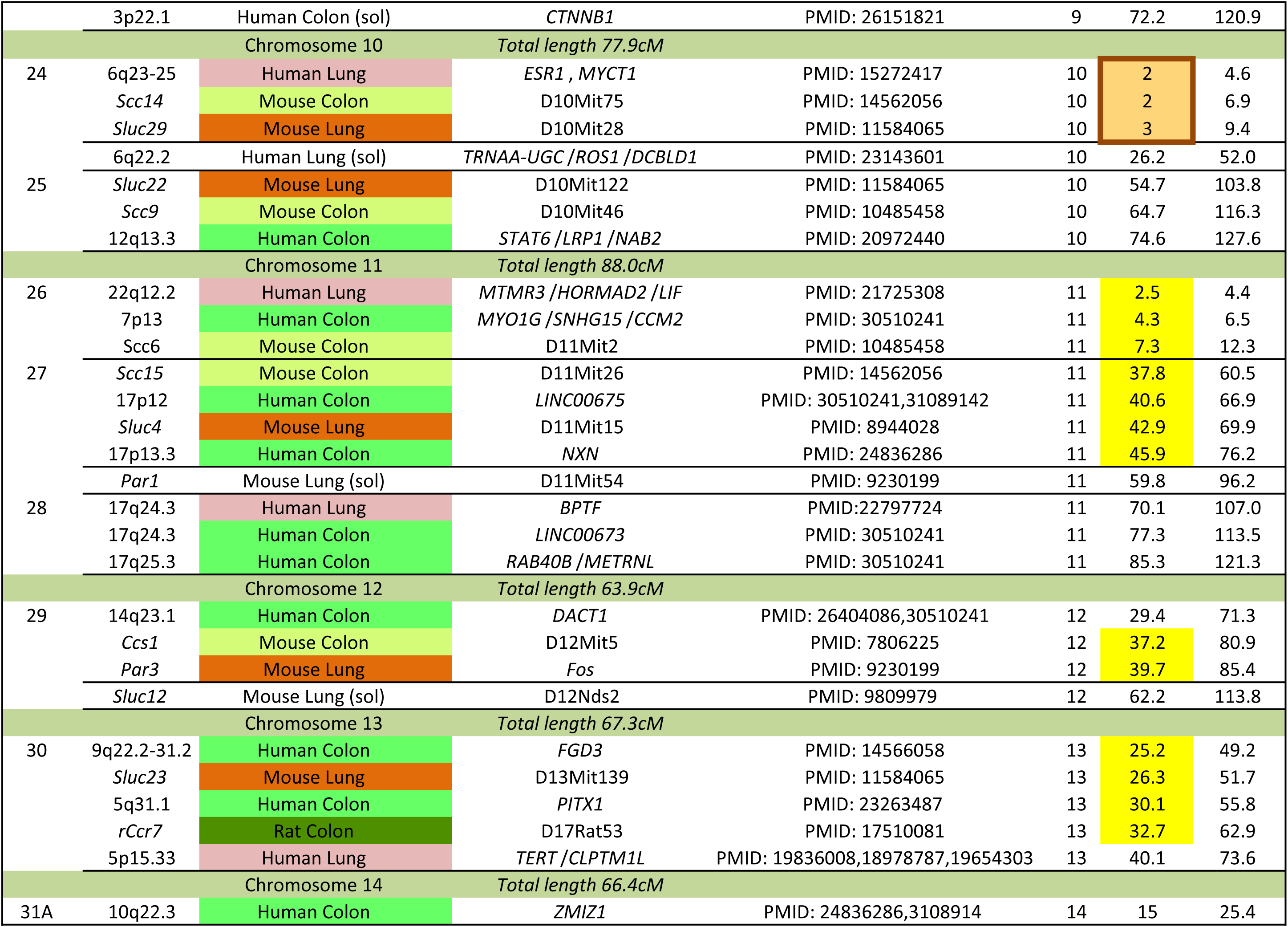
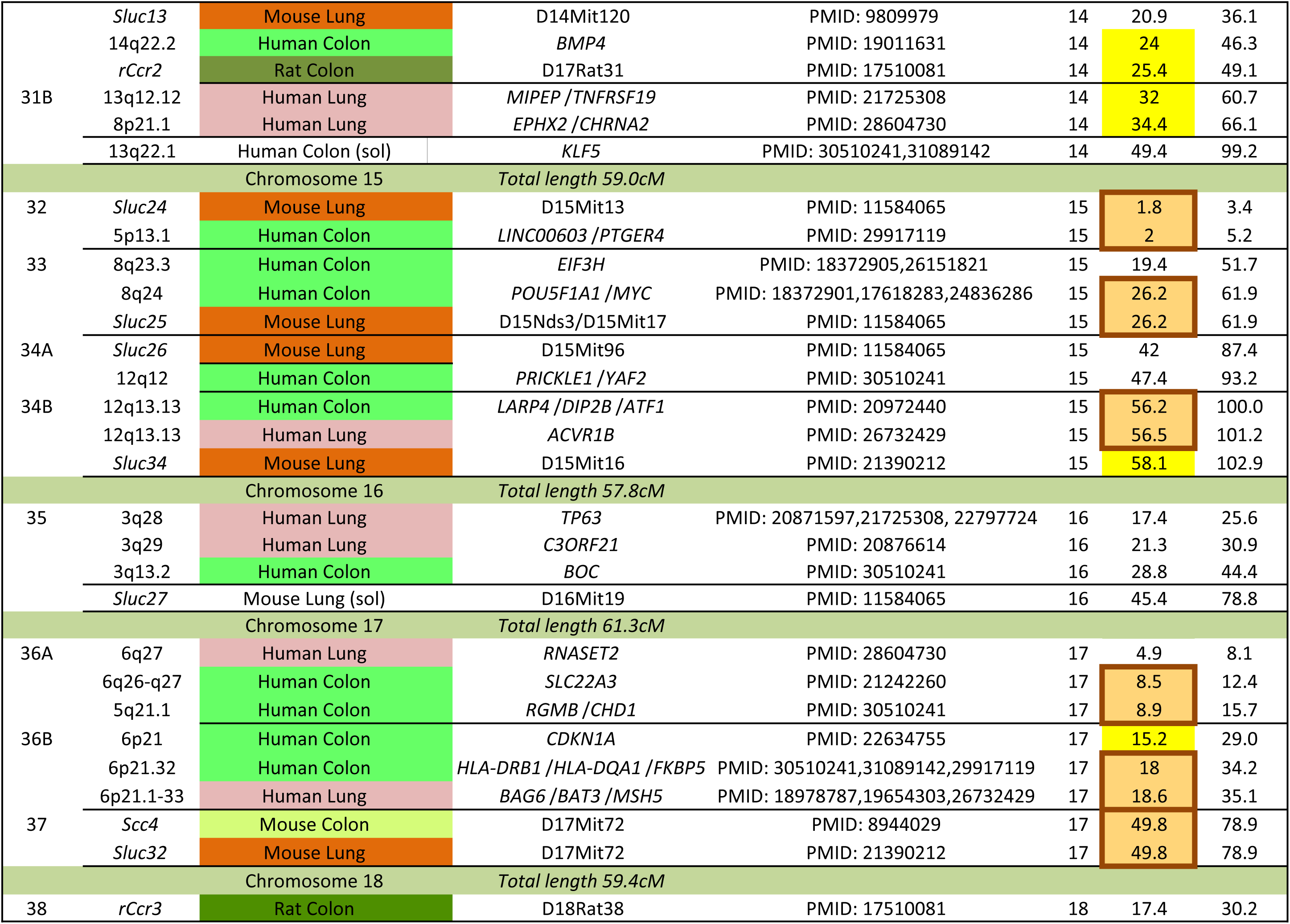
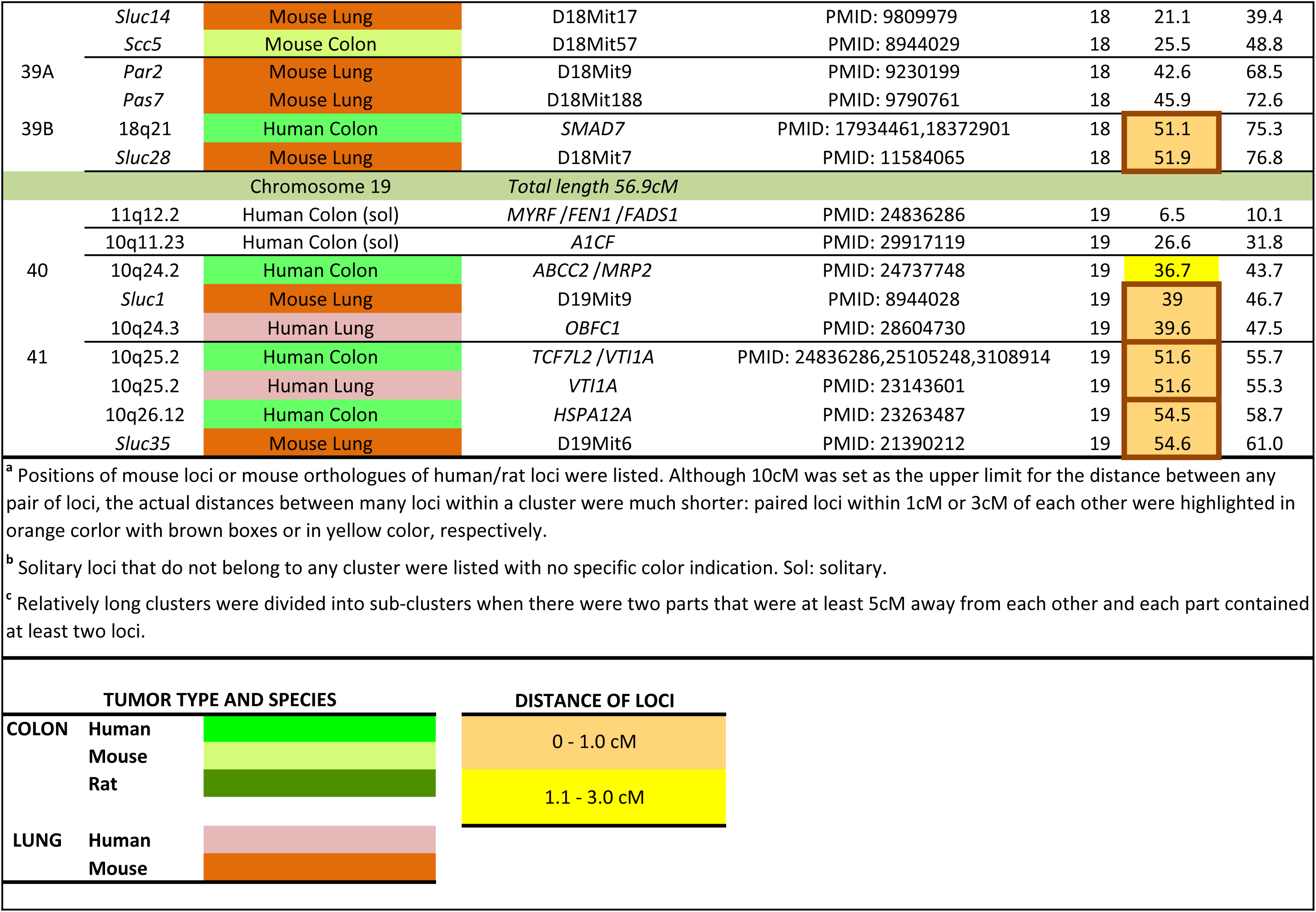
Co-localization of colon and lung cancer susceptibility loci identified in human, mouse and rat.

#### c. Interspecific conservation of linkage

The linkage data comprised 159 chromosomal locations of colon and lung cancer susceptibility loci. The orthologous mouse locations of 142 of them formed 41 clusters (Table 1) with the maximal distance of loci within a cluster being 10 cM. This conserved linkage involved 64 human colon cancer and 22 human lung cancer QTLs detected relatively early in the GWAS studies^15,14,4,11^. Such early detected loci appear to be more often replicated in subsequent GWAS studies^17^ whereas the cancer QTLs detected in later larger size GWAS studies exhibit progressively smaller effects^18^.

32 of the 41 clusters (78%) contained loci from at least two species (Table 1). Tests of colon tumor susceptibility in additional mouse strains and different mouse subspecies mapped 3 additional susceptibility loci into the clusters listed in Table 1: Scc22 - cluster 9, Scc25 - cluster 16, and Scc27 - cluster 38^7^.

Inter-organ clustering. 37 of the 41 clusters (90%) contained both colon and lung cancer susceptibility loci (Table 1). Among the 159 analyzed loci, 77 of 95 colon cancer loci co- localized with at least one lung cancer locus: 22 of them within <1 cM and 18 within 1-3 cM; 56 of 64 lung cancer loci co-localized with at least one colon cancer locus: 22 of them within <1 cM and 12 within 1-3 cM.

Density of the clusters. The most numerous clusters were very short: 54/159 loci were in clusters shorter than 1 cM, which included 10 clusters shorter than 0.1 cM, 8 clusters 0.1-0.5 cM long, and 7 clusters 0.5-1.0 cM long. Only 17 of the 159 loci did not belong to any cluster.

#### d. Statistical evaluation of clustering

Statistical analysis supported the dominance of extremely short clusters. Monte Carlo simulations using 100,000 simulated databases at window widths of 1, 2, 5, 10, 15, and 20 cM indicated that the p values of the deviation of the actual distribution of the colon and lung susceptibility loci from random distribution at these window widths were 0.00067, 0.093, 0.070, 0.056, 0.799, and 0.44, respectively. Poisson analysis shows that 142 colon and lung cancer susceptibility loci map into 41 clusters with a total length of 341 cM, whereas 17 non-clustered loci map into the remaining 1100 cM, which when compared with the total length of mouse autosomal genome of 1441 cM, indicated a p<0.001. In contrast, the expected average random distance of the 159 cancer susceptibility loci in the autosomal mouse genome (1441 cM long) was 9.06 cM.

#### e. Concordant RC strain susceptibility to colon and lung tumors

The colon tu^•^m^•^ or numbers^3,5,7,8^ as well as tumor load or lung tumor number^5,9,11^ were evaluated. We used four mouse RC strains, each containing a different random set of 12.5% genes of the strain STS (highly susceptible to colon tumors^3^) and 87.5% of genes of strain BALB/c (resistant to colon tumors^3^). We have shown in independent experiments that the colon tumor susceptible RC strains CcS-19 and CcS-11 have also significantly higher lung tumor load and lung tumor number than the colon tumor resistant strains CcS-20 and CcS-10, respectively (p<0.0001 and p<0.0012, Wilcoxon rank sum two sample test)^5^. The lung tumor numbers were determined in a separate experiment using F1 hybrids of these four CcS strains with FVB mice because of shortage of CcS mice. A comprehensive evaluation of the concordance of colon and lung tumor numbers of these RC strains was performed by t-distributed linear contrast test, which indicated that the strains CcS-10 and CcS-20 differ. significantly in lung tumor numbers from the strains CcS-11 and CcS-19 (p = 0.0005, 44 degrees of freedom).

### 2. Systemic linkage of cancer susceptibility genes with the genes regulating protein phosphorylation

Changes in protein phosphorylation are critical components of tumorigenesis. They are regulated among others by a number of protein tyrosine phosphatases^19^ and driver protein kinases^20^ belonging to several structurally and functionally distinct groups.

#### a. Protein tyrosine phosphatases

Protein tyrosine phosphatase superfamily contains enzymes that remove phosphate groups either strictly from phosphorylated tyrosine, or also from phosphorylated serine and threonine. We tested the linkage of colon/lung cancer susceptibility loci with enzymes of the largest, type-I cysteine-based superfamily. It contains two large families and three additional classes of enzymes^19,21,22^. One family contains 21 receptor and 17 non-receptor classical protein-tyrosine phosphatase enzymes with a strict tyrosine specificity, the second a group of seven subfamilies comprising 61 dual specificity phosphatases (DUSP), which besides tyrosine can dephosphorylate also serine and threonine. Finally, this superfamily also contains three separate classes of phosphatases with one, three, and four enzymes, respectively.

We tested the linkage of type-I cysteine-based superfamily phosphatases with colon/lung tumor susceptibility loci listed in Table 1. The results in Table 2A show in the left half the map position of the individual protein tyrosine phosphatases and the nearest colon/lung cancer susceptibility loci. In the right half are the linked pairs of phosphatase and colon or lung cancer susceptibility loci ranked by their distance. This provided very strong evidence of linkage. Of the 107 phosphatase genes were 25 (23.4%) located within 1 cM or less of a colon/lung tumor susceptibility locus, 27 (25.2%) within 1.1-3 cM (total 48.6%, Poisson test p value 1.11cx10^-16^) and 16 (15.0%) within 3.1-5 cM of a colon/lung tumor susceptibility locus. Only 2 (1.9%) were at 8.1-10 cM apart and 23 (21.5%) were non-linked, i.e., separated by 10 or more cM. The average distance between a phosphatase gene and a linked colon/lung tumor susceptibility locus (i.e., distant maximally 10 cM) is 2.73 cM, median 2.15 cM. Testing separately the linkage of classical tyrosine phosphatases and of the DUSP phosphatases provided similar results in the two large groups, the average and median distance from the nearest colon/lung susceptibility locus was 3.21 and 2.45 cM for classical phosphatases, and 2.37 and 2.00 cM in DUSP phosphatases. We did not test linkage of colon/lung cancer susceptibility loci with the serine-threonine phosphatases, because their substrate specificity in many instances is determined by combination of non-linked genes.

**Table 2A.**
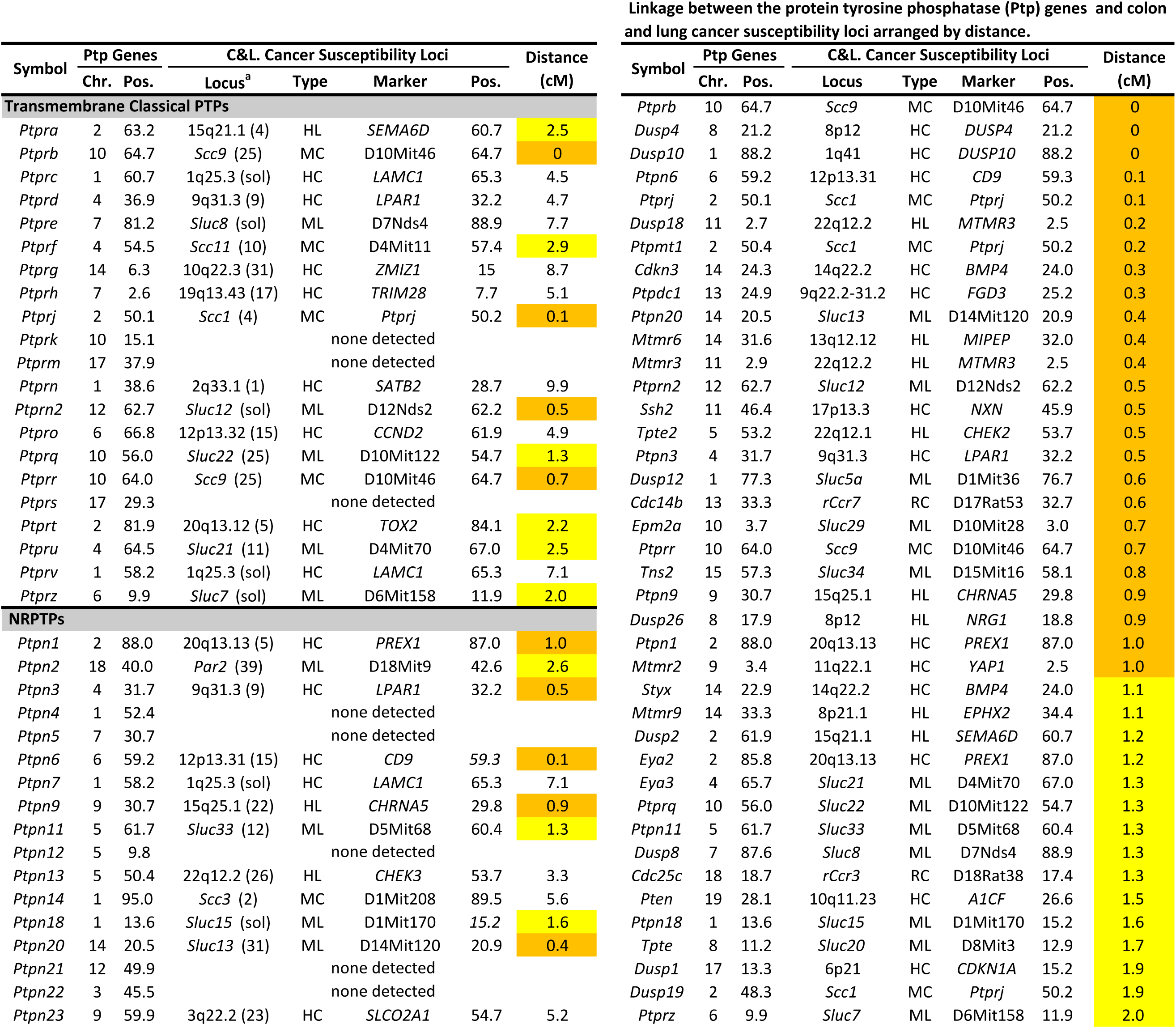
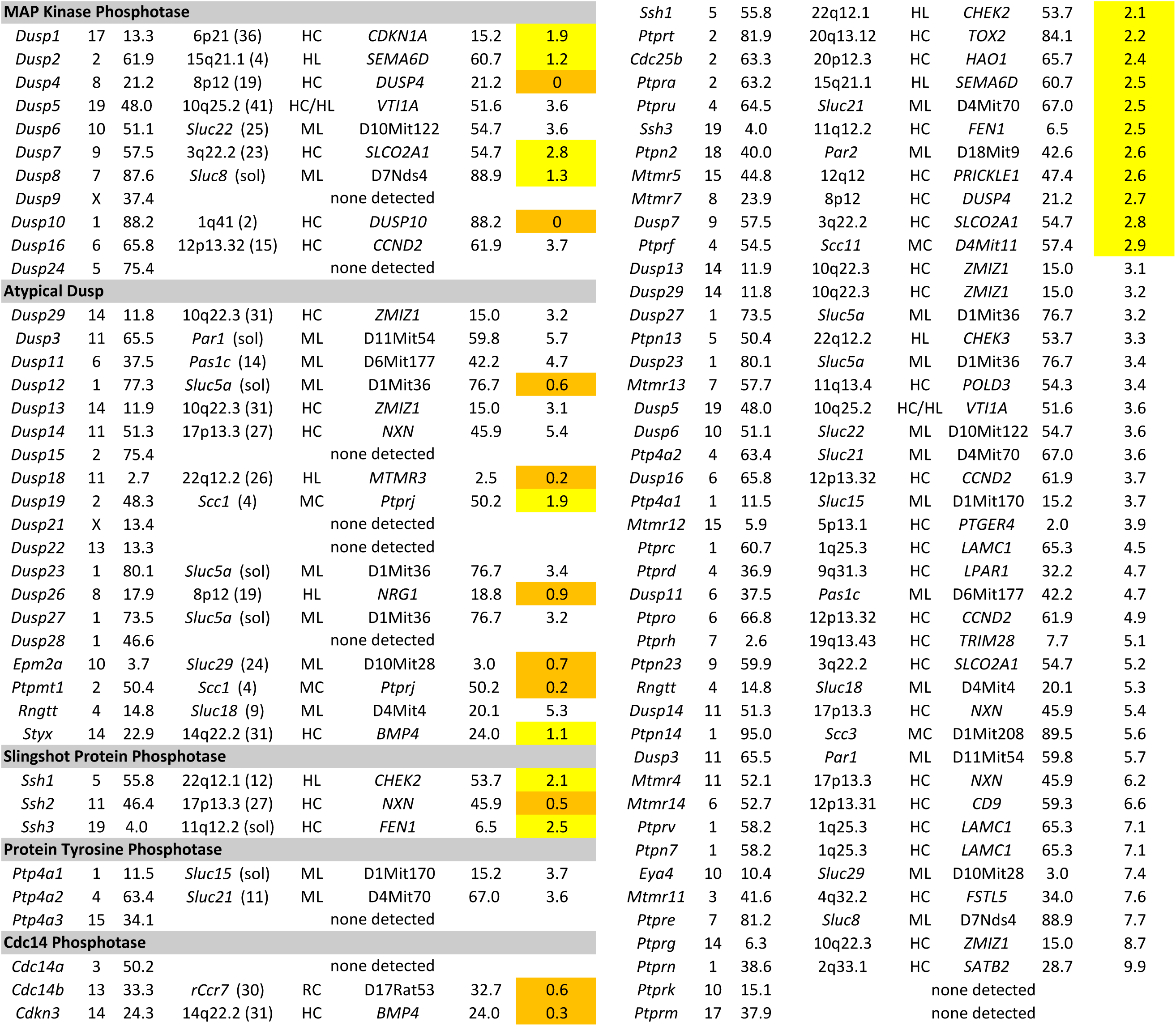
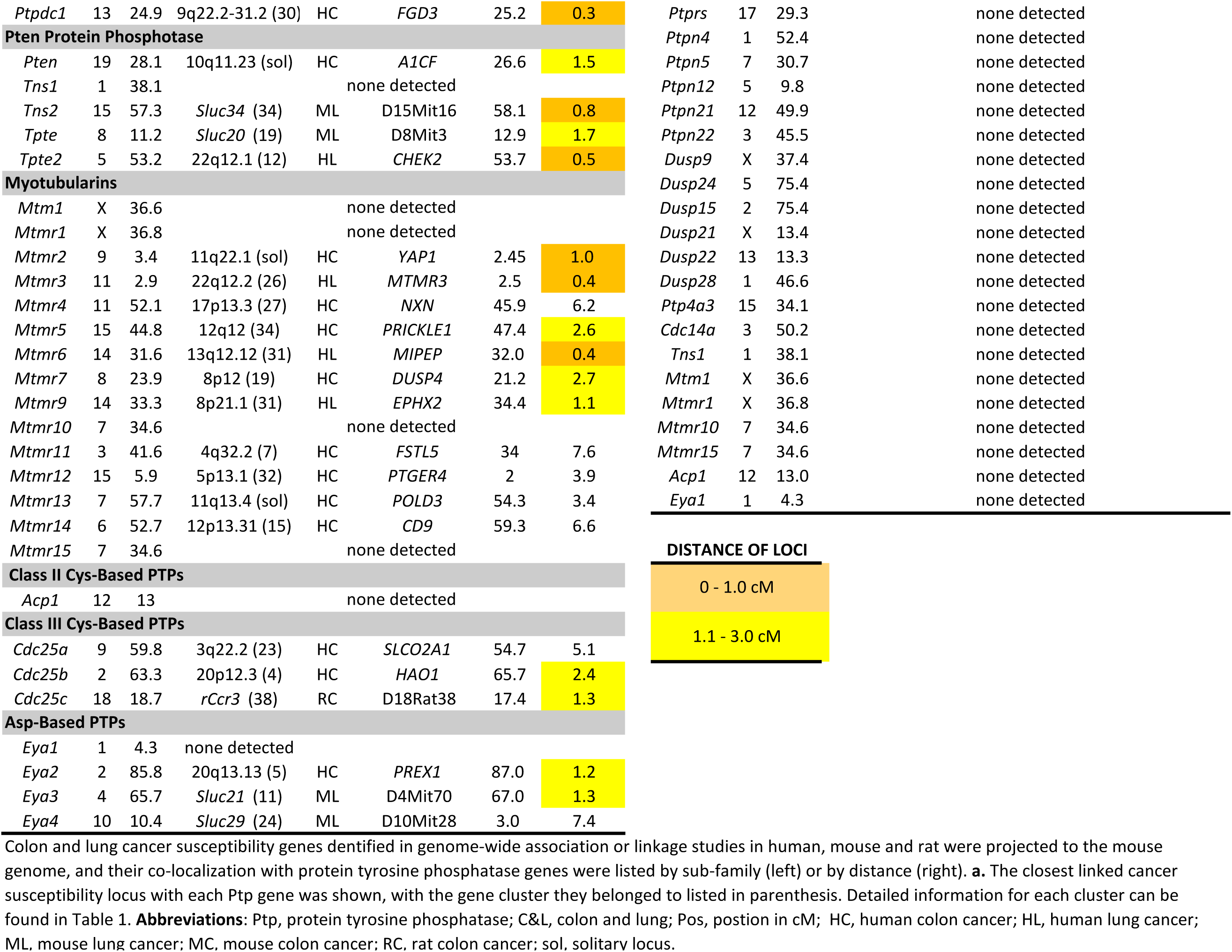
Linkage between the protein tyrosine phosphatase (Ptp) gene family and colon and lung cancer susceptibility loci.

#### b. Protein kinases

We tested linkage of colon/lung cancer susceptibility loci with the class of cancer driver protein kinases listed in^20^. The orthologous positions of these human genes on mouse chromosome map were compared with those of colon and lung tumor susceptibility loci of the three species listed Table 1. This data (Table 2B) was organized in the same way as in Table 2A and revealed that 48 of 56 (85.7%) protein kinases are linked to a colon or lung cancer susceptibility gene, 17 of them at a distance of 1 cM or less (30.4%), and 14 at a distance of 1.1-3 cM (22.0%), (total 52.4%, Poisson test p value <1.0 x 10^-30^), 6 at a distance of 3.1-5cM (10.7%). Only 4 (5.4%) were distant 8.1-10 cM and 8 (14.3%) were separated by 10 or more cM (19.6%). Their average distance from the nearest colon or lung cancer susceptibility locus was 2.79 cM, median 2.0 cM.

**Table 2B.**
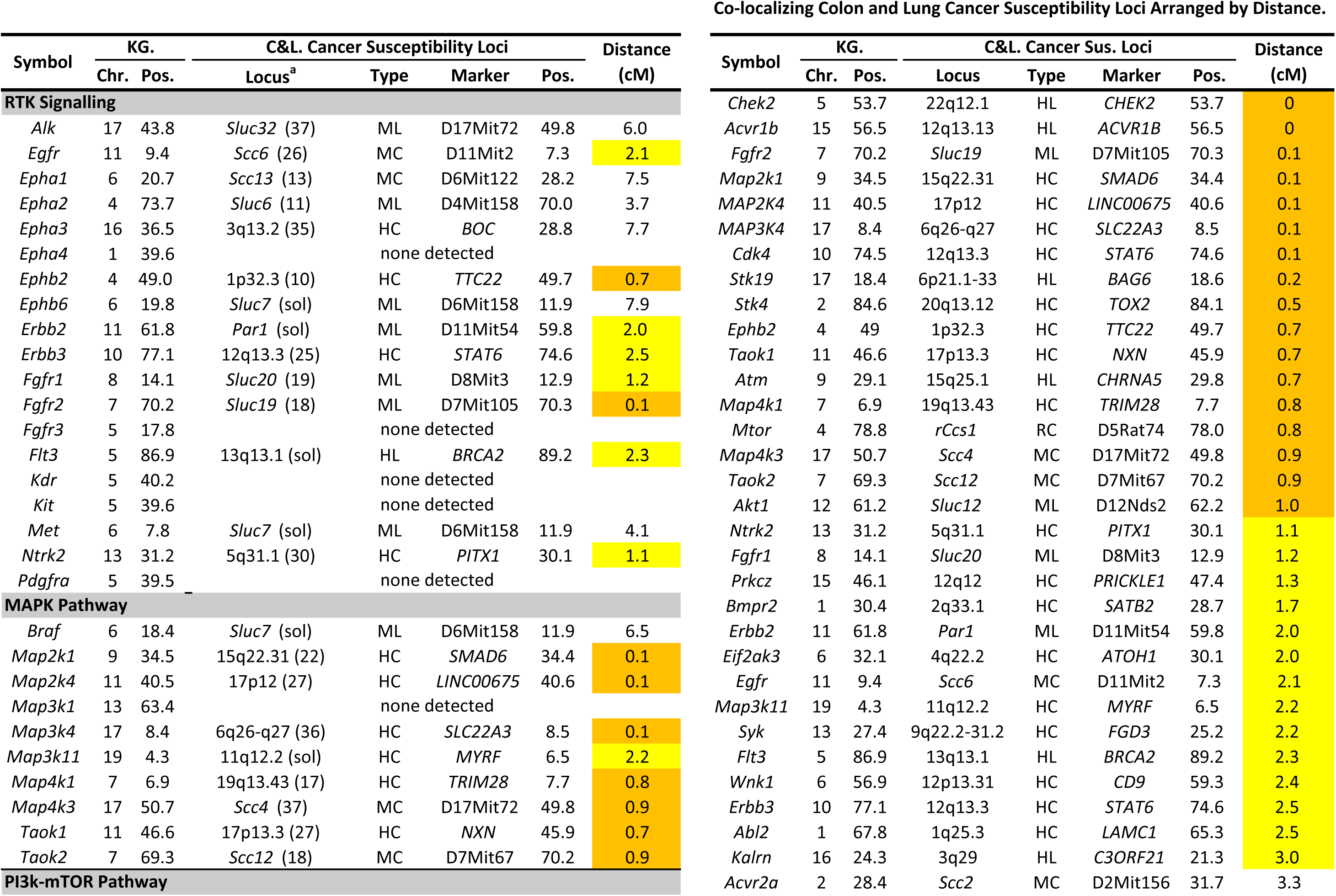
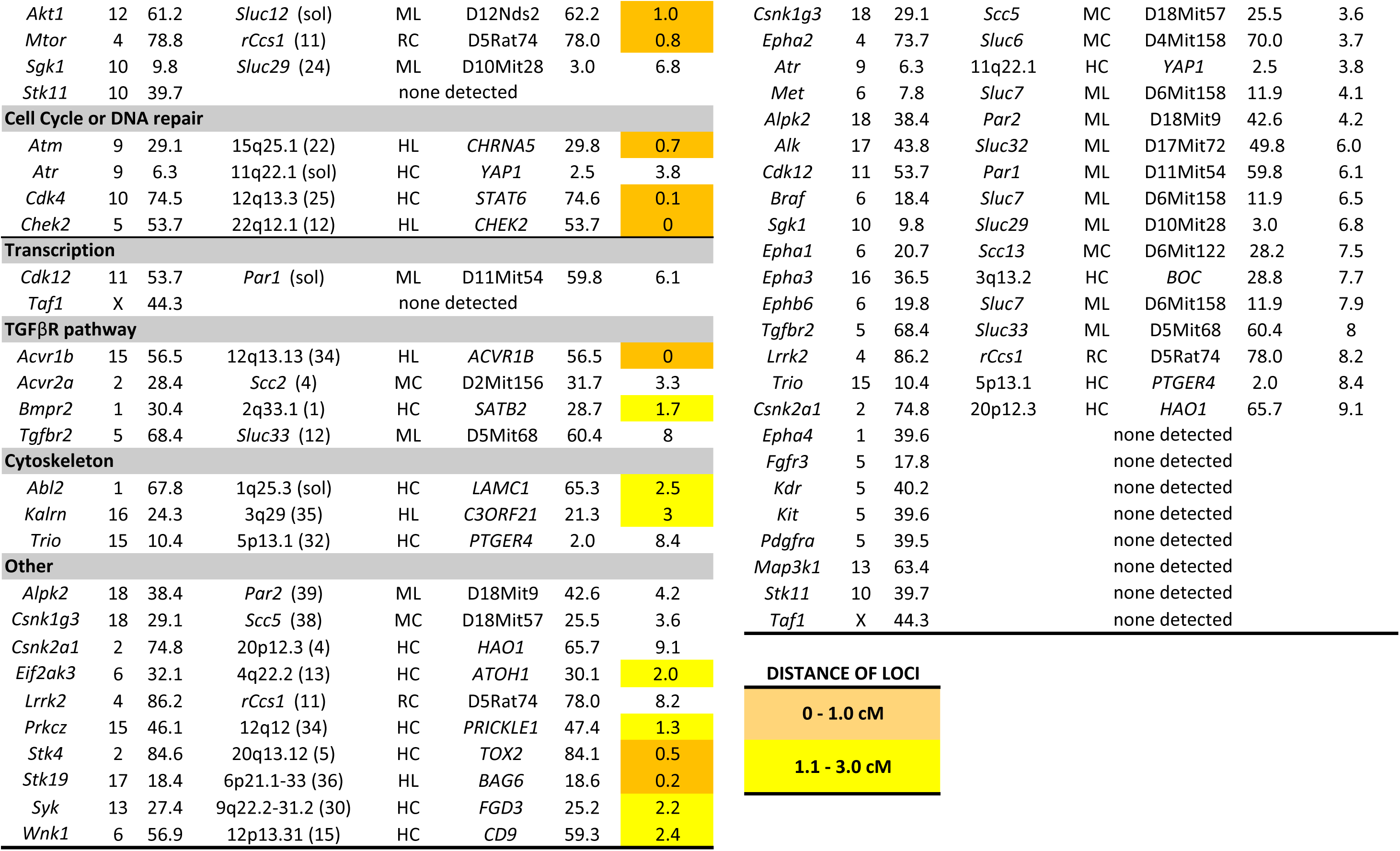
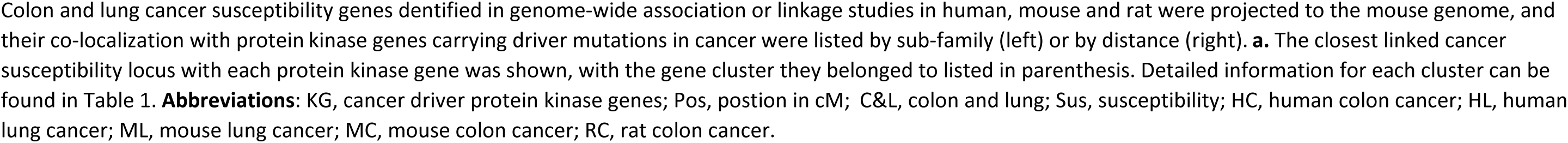
Linkage between cancer driver protein kinase genes and colon and lung cancer susceptibility genes.

#### c. Linkage of protein tyrosine phosphatase and protein kinase genes to susceptibility loci to other cancers in humans

We next tested the linkage of these two groups of genes involved in protein phosphorylation and dephosphorylation with cancer susceptibility in humans, using the data of Sud et al^4^. These early GWAS data on 29 human cancers comprising 962 GWAS defined susceptibility loci at 263 non-redundant chromosomal locations. They demonstrated linkages of protein tyrosine phosphatases with cancer susceptibility genes in 94 out of 107 cases (Table 3) and driver protein kinases in 50 out of 56 cases (Table 4). Poisson analysis using an estimated length of 3055 Mb of human genome revealed highly significant aggregation of protein tyrosine phosphatase genes, protein kinase genes, or both combined, in genomic regions containing susceptibility loci for various types of cancer. For protein tyrosine phosphatase genes, 28 (26.2%) were at 1 Mb or less (Poisson test p value <3.2 x 10^-7^), 53 (49.5%) at 3 Mb or less (Poisson test p value <5.6 x 10^-6^), and 16 at distance 3.1-5 Mb (15.0%). Only 13 (12.1%) were separated by 10 or more Mb. Their average distance from the nearest cancer susceptibility locus was 3.0 Mb, median 2.0 Mb.

**Table 3.**
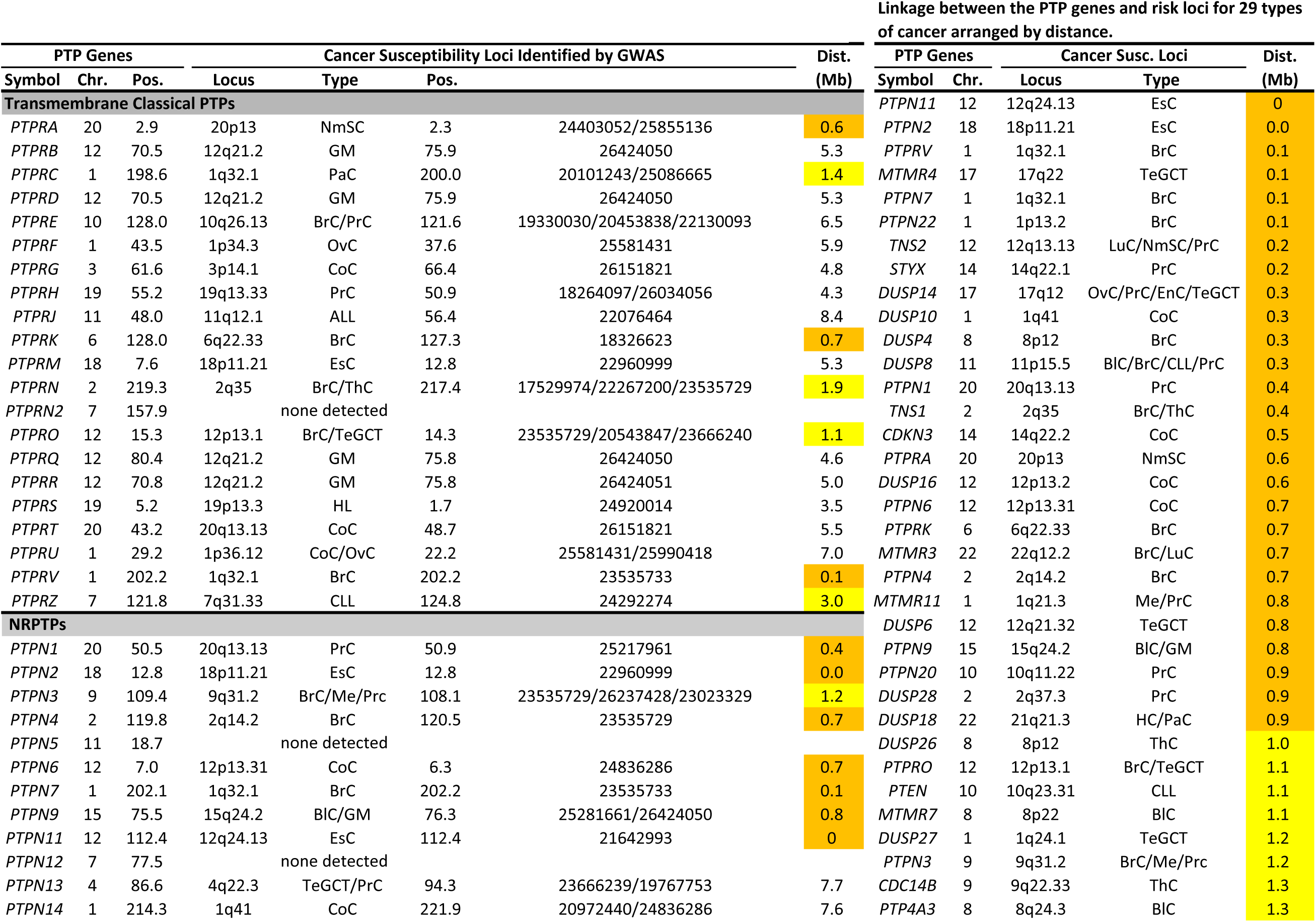
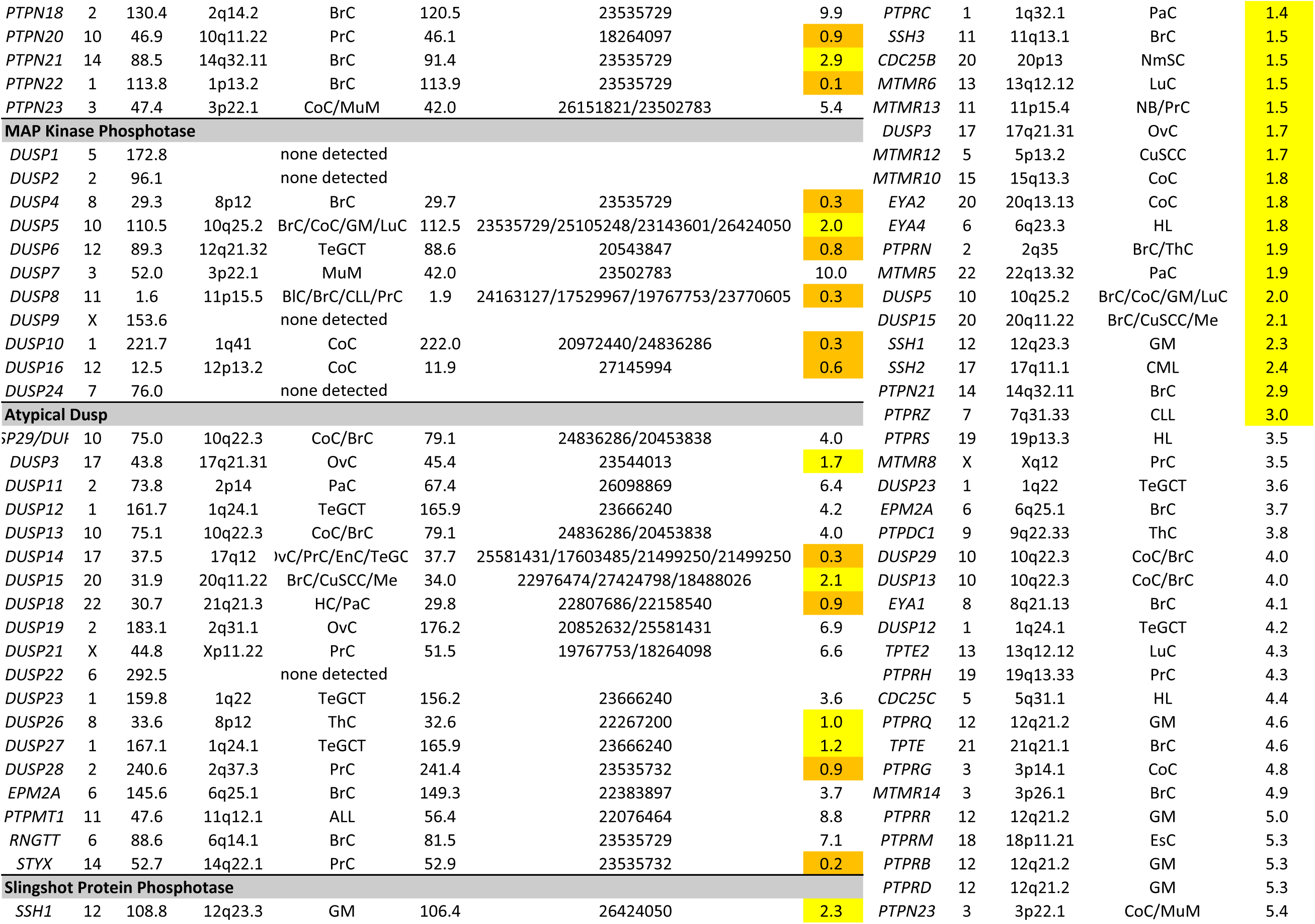
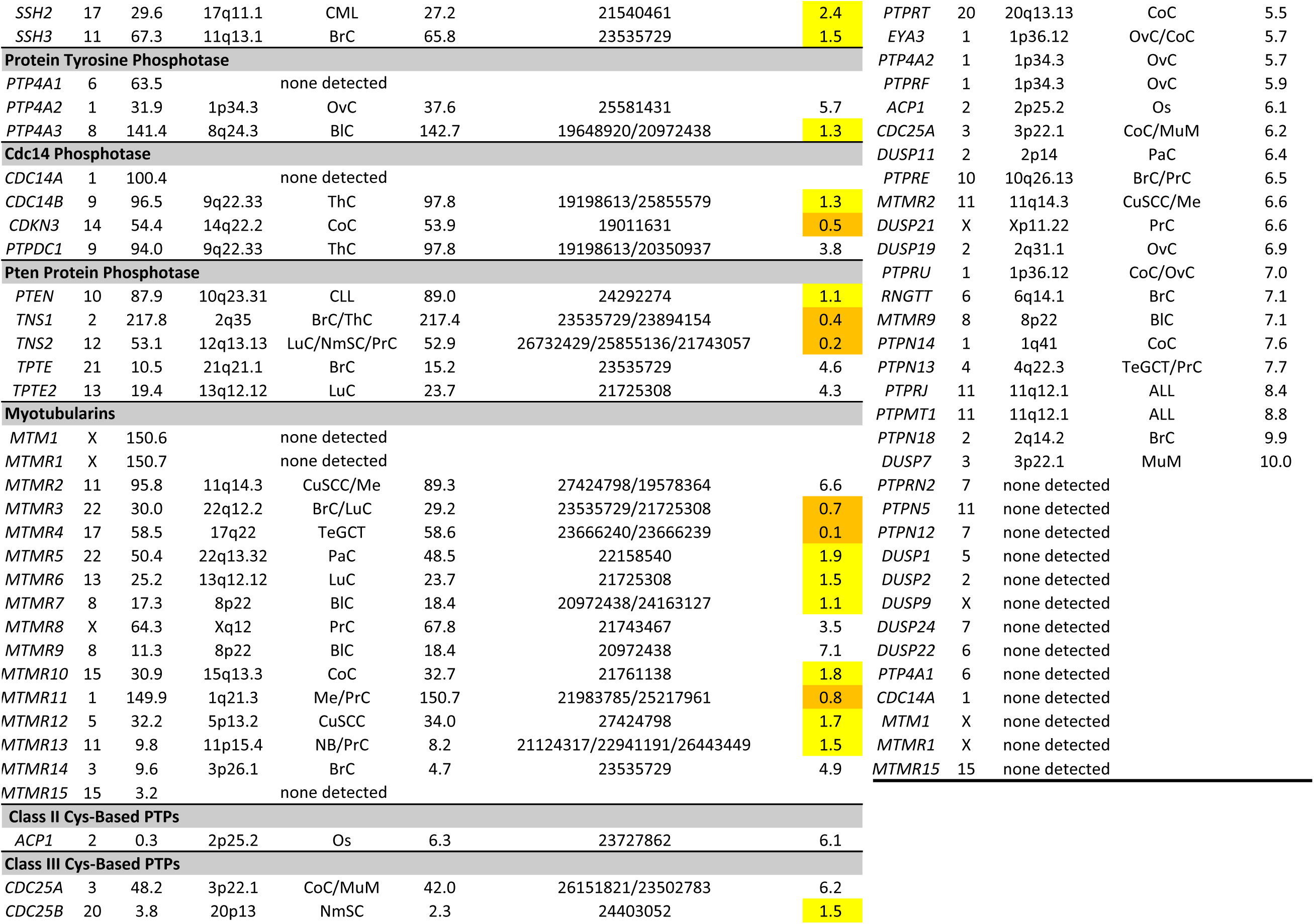
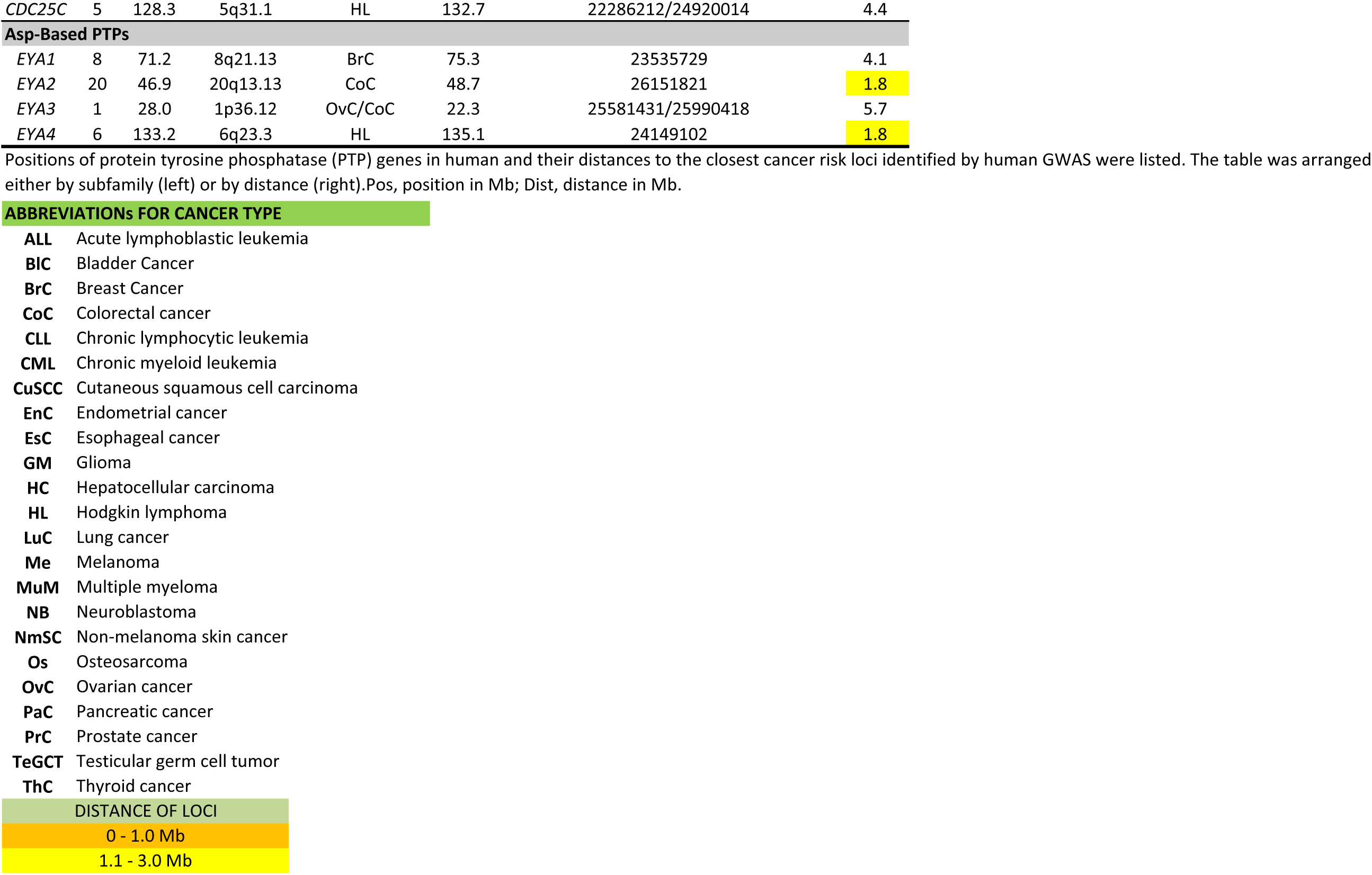
Linkage between the protein tyrosine phosphatase (PTP) gene family and risk loci for 29 types of cancer identified by GWAS in human.

**Table 4.**
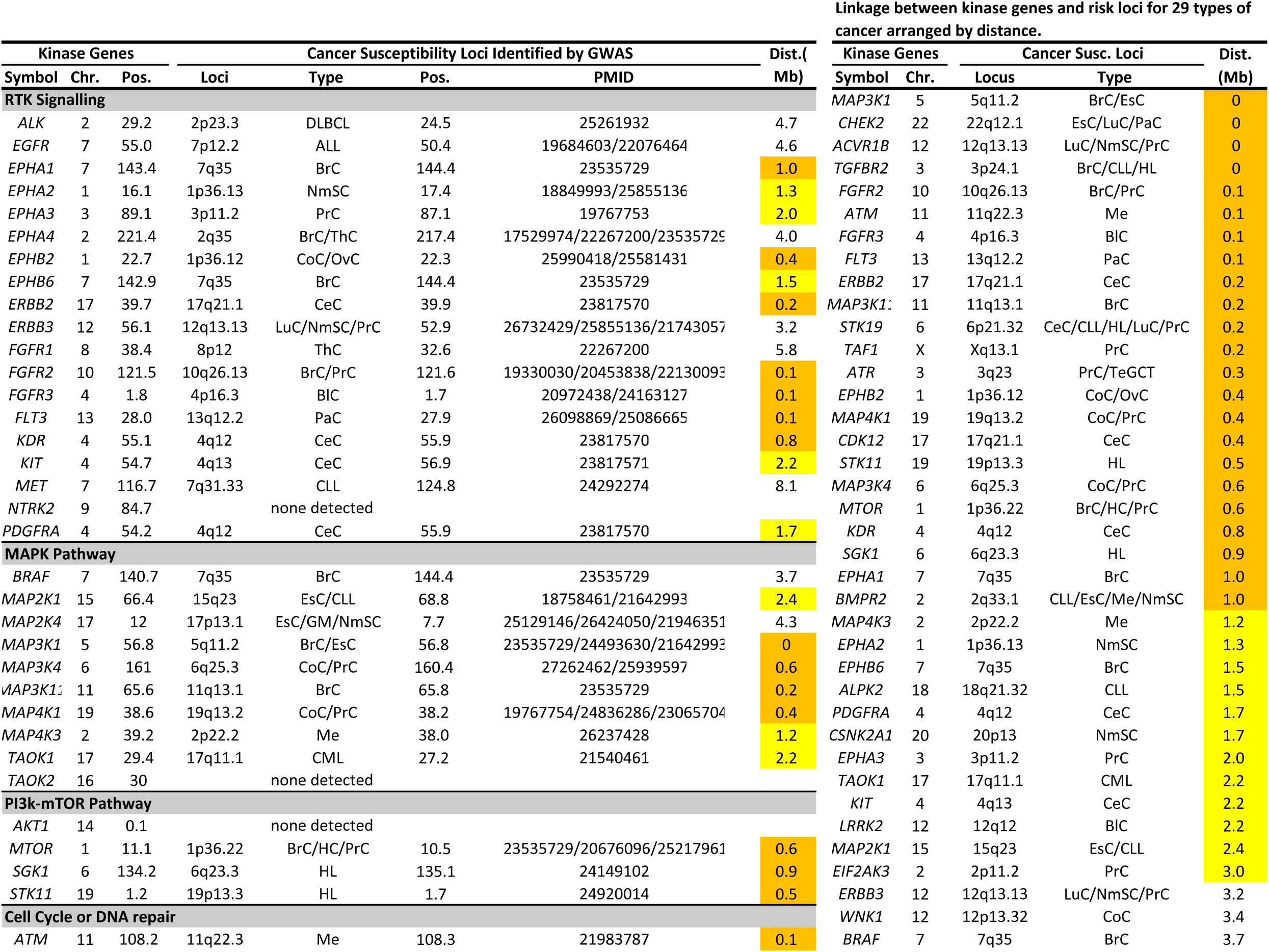
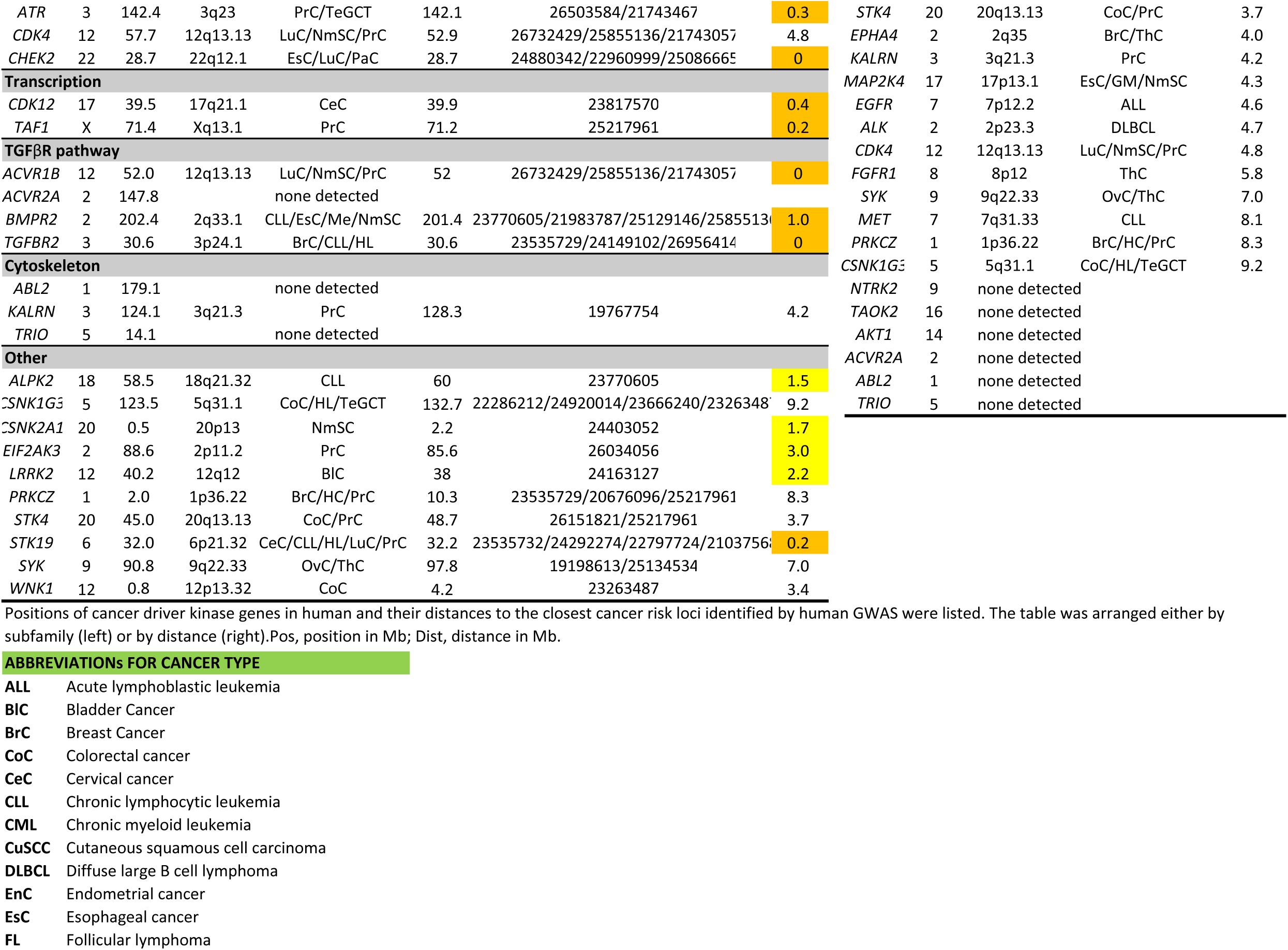
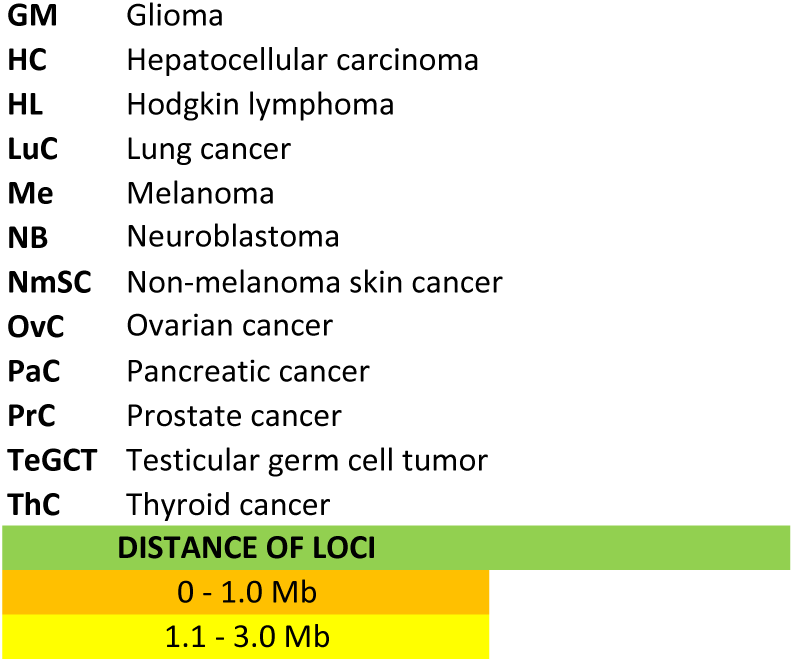
Linkage between the cancer driver protein kinase genes and risk loci for 29 types of cancer identified by GWAS in human.

Similarly for protein kinase genes, 23 (41.1%) were at 1 Mb or less (Poisson test p value <1.5 x 10^-9^), 35 (62.5%) at 3 Mb or less (Poisson test p value <1.9 x 10^-6^), and 10 at a distance of 3.1- 5 Mb (17.9%). Only 6 (10.7%) were separated by 10 or more Mb. Overall, their average distance from the nearest cancer susceptibility locus was 2.2 Mb, median 1.4 Mb (Table 4). To rule out any potential bias rendered by colon cancer and lung cancer, we removed susceptibility loci for these two types of cancer from the analysis and repeated the Poisson analysis. The association of protein tyrosine phosphatases (3.1x10^-6^ and 4.8x10^-5^) for distance of 1 and 3 Mb and protein kinases (1.8x10^-8^ and 2.4x10^-6^) for distance 1 and 3 Mb with cancer susceptibility loci remained significant.

## Discussion

The colon and lung cancer susceptibility genes, which have been generally considered different and independent entities spread randomly over the genome, are according to our results in fact evolutionarily conserved and located in a set of relatively short genomic regions which also contain most protein kinases and protein tyrosine phosphatases.

### 1. Shared characteristics of colon and lung cancer susceptibility

While the molecular biology of colon and lung tumors are different, our data indicate that they share important genetic characteristics. Their pairwise chromosomal co-localization in different species has been conserved for >70 million years, and they are concentrated in several relatively short clusters on different chromosomes that collectively represent a limited part of the genome. Their pairs are often associated with closely linked SNP markers and the linked pairs of colon and lung susceptibility genes tend to exhibit a correlated phenotype (either both susceptible or both resistant). There are several examples of inter-species correlations of colon and lung cancer susceptibility. Three suggestive loci affecting more than one human cancer^23^ fall unto colon-lung loci 1p36.3 10q26.13, 10q26.12 and two pairs of colon and lung cancer loci^24^ are confirmed in Table 1: one at 19q13.2 is part of cluster 17 on chromosome 7 and one at 20q13.33 is part of cluster 6 on chromosome 1, which contains also the independently identified mouse lung tumor susceptibility locus Sluc17^3^ (Table 1). However, the considerable linkage and overlap of colon and lung cancer susceptibility genes and suggested correlation of their individual effect are not reflected in a correlation of the clinical incidences of these two cancers. Therefore, the multiple common characteristics shared by colon and lung susceptibility genes require mechanistic studies.

### 2. Linkage with protein tyrosine phosphatases and protein kinases

The two different groups of proteins, protein tyrosine phosphatases^19,21,22^ and cancer driver protein kinases^20^ are both significantly linked with colon/lung tumor susceptibility loci (Table 2A, 2B). The linkages of protein kinases with colon/lung cancer susceptibility loci are approximately as close as those of protein tyrosine phosphatases genes but seem to involve mostly different colon/lung cancer susceptibility loci. Both protein tyrosine phosphatases and cancer driver protein kinases are involved in cancer development. Although originally protein tyrosine phosphatases were considered tumor suppressors, they can be involved both in suppression as well as in cell transformation^25–28^. This linkage also includes cancer susceptibility genes in humans, including other than colon and lung (Tables 3 and 4). The expansion of possibilities of proteomics-based analysis of protein kinases and protein tyrosine phosphatases^28,29^ can improve the definition of action of these genes. The protein kinases analyzed in this study belong to several families of protein kinases and one of their shared features is their driver function in various cancers, not only of colon and lung^20^.

### 3. Potential implications

Colon and lung cancer are important causes of cancer morbidity and mortality. Previously many colon and lung cancer susceptibility loci in mouse and human have been identified with specific genes, including oncogenes and tumor suppressor genes, as well as several protein phosphatases and kinases. These genes are included in Table 1 and their possible relation to the novel characteristics described here require future elucidation. Here we discussed two unexpected new phenomena associated with colon/lung cancer susceptibility genes: the extensive evolutionary conserved concordance of their pair-wise linkage and concordant impact of their linked pairs on tumorigenesis in the two organs, as well as and their unexpected close linkage with protein tyrosine phosphatases and cancer driver protein kinases. They suggest a possible presently unknown but related genetic mechanisms that may modulate the tumorigenesis. They also raise the question about the possible functional heterogeneity of distinct chromosomal segments carrying susceptibility genes as well as protein tyrosine phosphatases and protein kinases.

A future insight into nature of this control can be obtained by comparing global gene expression^29–31^ in colon and lung tissues and tumors of RC strains with pronounced concordant colon/lung tumor susceptibility or resistance. It could indicate the expression patterns that are organ specific and are related to degree of susceptibility and detect the possible trans-regulatory effects of STS-derived genes on BALB/c-derived genes. Finally, a study of 3D chromatin organization^32^ in the evolutionary conserved regions containing cancer susceptibility genes might indicate the extent of their conservation and diversity.

Importantly, we observed in addition to colon and lung cancer susceptibility genes the same type of linkage to protein tyrosine phosphatases and protein kinases in susceptibility genes to other cancers. This included linkage to a protein tyrosine phosphatase in 94 out of 107 of cancer susceptibility loci and linkage to protein kinases in 50 out of 56 cancer susceptibility loci. The biological significance of these linkages remains to be investigated. Potentially it can change the current view of cancer susceptibility genes from largely organ-specific solitary genes associated individually with oncogenes and tumor suppressor genes, into an integrated system of several types of genes including evolutionarily conserved multi-organ specific genes associated with oncogenes, tumor suppressor genes, and simultaneously also with several classes of protein tyrosine phosphatases and protein kinases. The mouse recombinant congenic strains can be particularly suitable to characterize its multimodal structure in relation to multistep tumorigenesis.

In summary, the colon and lung cancer susceptibility genes that have been considered biologically and genetically non-related entities share across organ and species boundaries several common features: pair-wise genetic co-localization conserved over 70 million years, linkage with protein tyrosine phosphatases and cancer-driver protein kinases, and concordant effect on tumorigenesis in the two organs. Functional exploration of this cross-organ and cross-species relatedness may provide a more comprehensive and systemic understanding of genetic components of carcinogenesis.

## Methods

### Ethics statement

All animal experiments were approved by the IACUC committee at Roswell Park Comprehensive Cancer Center (IACUC protocol 9O5M).

### Mice

We used 20 BALB/c-c-STS (abbrev. CcS) RC strains (CcS-•1· through CcS-2O), each of which inherited a random part of 87.5% of the genome from the “background” BALB/c strain and 12.5% of the genome from the “donor” STS strain^3,5^ Mice received acidified drinking water (pH 2.5-3.0) and a standard laboratory diet (LM-485, Harlan Teklad, U.S.) ad libitum and at the end of experiment were euthanized by CO2 asphyxia.

### Tumor induction and histological analysis

<u>Lung tumor induction</u>: Pregnant female mice were given one intraperitoneal (i.p.) injection of 30 mg/kg body weight of the carcinogen N-ethyl-N-nitrosourea (ENU, Sigma N3385) dissolved in phosphate-buffered citric acid (pH 5.8) at day 17 of gestation. The offspring of carcinogen-injected females were thus exposed to ENU transplacentally. At the age of four months the lungs were removed, embedded in histowax, sectioned semi-serially and evaluated for tumors. <u>Colon tumors</u> were induced in adult mice with eight weekly subcutaneous injection of 15 mg/kg body weight of azoxymethane (AOM, Sigma A5486). Mice were euthanized four months after the last injection and their colons removed, flushed, longitudinally cut, splayed open, and examined for numbers and locations of tumors.

### Genotyping

The positions and lengths of the most ‘donor’ strain-derived chromosomal regions in CcS RC strains have been determined with more than 500,000 single nucleotide polymorphism (SNP) markers at the Jackson Laboratory and multiple microsatellite markers across the whole genome.

### Definition of clusters of loci

Susceptibility loci for colon and lung cancers were retrieved from literature and the NHGRI GWAS catalog (for human GWAS). Only loci with a genome-wide significance in humans were used (p<5x10^-8^ for GWAS) or P<0.05 after Bonferroni multiple testing correction for loci in rat and mouse. Loci identified in human and rat were projected onto the mouse genome based on positions of their orthologous regions in mice, acquired from the major genome browsers, i.e. the Jackson Laboratory (https://informatics.jax.org), the Rat Genome Database (https://rgd.mcw.edu), Ensemble (https://www.ensembl.org) and the UCSC genome browser (https://www.genome.ucsc.edu). We set 10cM as the upper limit for the distance between paired loci within a cluster, but the actual distances between most loci in a cluster were much shorter (Table 1).

### Linear contrast analysis of the lung tumor count data

Data consist of tumor counts as outcome variable, with listing of strain (CcS-10, CcS-11, CcS-19, and CcS-20) and induced tumor type (lung). The analysis aimed to examine the specific contrast of testing for a difference between the strains (CcS-10, and CcS-20) and the strains (CcS-11 and CcS-19), among the lung cancer induced mice. The results from testing this contrast are given here: > summary(G, test=adjusted(’single-step’))Simultaneous Tests for

### General Linear Hypotheses

Multiple Comparisons of Means: User-defined Contrasts Fit: lm(formula = TumorCts - Strain, data= lungdat) Linear Hypotheses: cl0c20vscllcl9 == 0, Estimate= -6.470, Standard error 1.718, t value= -3.765, Pr(>JtJ) 0.000491 *** (44 degrees of freedom)

(Adjusted p values reported -- single-step method)

These results indicate that there is a significant value for the contrast, which would be interpreted as demonstrating that, among the mice treated to induce lung tumors, the two strains CcS-10 and CcS-20 differ significantly in tumor counts compared to the two strains CcS-11 and CcS-19. These aren’t pairwise comparisons, but rather using the contrast formulation to aggregate the strains of interest to examine for significant differences in tumor counts.

### Assessment of colon and lung tumor clustering

Monte Carlo multi-window analysis was used to assess the distribution of positions of the mouse colon and lung cancer susceptibility loci in mouse autosomal genome and the orthologous positions of human and rat colon and lungcancer susceptibility loci projected on the mouse autosomal genome as listed in Table 1. We accounted for chromosome length, and only considered within-chromosome chromosome distances. The test involved six window widths (1, 2, 5, 10, 15, and 20 cM) and 100.000 simulations for each of them.

### Poisson analysis

The degree of linkage of protein tyrosine phosphatases and protein kinases with colon/lung cancer susceptibility genes was assessed by the Poisson analysis of the proportion of the protein tyrosine phosphatase genes or protein kinase genes mapping to a colon or a lung tumor susceptibility locus to a distance 3 cM or smaller, respectively (calculated by the POISSON.DIST function in Excel).

## Acknowledgements

We thank Dr. Miranda Lynch from the Hauptman-Woodward Medical Research Institute in Buffalo, New York, for performing Monte Carlo simulations and linear contrast analysis, Dr. Alan Hutson, Chairperson of Department of Biostatistic and Bioinformatics of the Roswell Park Comprehensive Cancer Center for generous support during this work, and Dr. Marie Lipoldova, Institute of Molecular Genetics of the Czech Academy of Sciences in Prague for useful comments.

## Author information

Roswell Park Comprehensive Cancer Center, Division of Cellular and Molecular Biology, Buffalo, 14260 USA

Peter Demant

Tjanjin Key Laboratory of Exercise Physiology and Sport Medicine, Institute for Exercise and Health Science, Tianjin University of Sport, Tianjin, China

Lei Quan

Correspondence:

Peter Demant

This work had been supported by the NCI grant R01 CA 115158 and an institutional grant from the Roswell Park Comprehensive Cancer Center to P. Demant, and by NCI grant P30CA016056 to the Roswell Park Comprehensive Cancer Center supporting the use of its Biostatistics and Genomic Shared Resources and by the grant 16JCYBJC from the Natural Science Foundation from the Tianjin City and Tianjin Thousand Young Talents Plan to L. Quan.

